# *Candida albicans* Snf2 modulates the response to DNA damage by regulating gene expression and uptake of the genotoxic stressors

**DOI:** 10.64898/2026.02.13.705856

**Authors:** Sanju Barik, Rajan Kushwaha, Akansha Arora, Ketki Patne, Anusha Ghosh, Rohini Muthuswami

## Abstract

The SWI/SNF complex comprising of the catalytic subunit, Snf2, is a key regulator of gene expression and DNA damage repair in eukaryotic cell. *Candida albicans* Snf2 is known to regulate hyphal formation. In this paper, we have investigated the role of this protein in DNA damage response. We show that *Ca*Snf2 is required for cell division as deletion of both copies of *SNF2* leads to increased duplication time. The mutant cells form clumps with increased chitin and β-glucan deposition on the cell wall. The altered cell wall phenotype leads to reduced uptake of genotoxic stressors leading to increased resistance to both methyl methane sulfonate (MMS) and hydroxyurea (HU). In addition, resistance of *Casnf2Δ* cells to MMS also appears to be mediated by upregulation of *CaRAD9* expression by *Ca*Fun30, an ATP-dependent chromatin remodeling protein, and *Ca*Rtt109, a fungal-specific histone acetyltransferase. The response of *Casnf2Δ* to genotoxic stressors is at variance with the response of *Scsnf2Δ* mutant, highlighting the differences in DNA damage response/repair pathway between the two organisms. Finally, we show that *Casnf2Δ* mutants are extremely sensitive to azoles due to downregulation of multi-drug resistance pumps leading to reduced efflux of the drug.

## INTRODUCTION

*Candida albicans* is a commensal opportunistic fungus that causes candidiasis in immunocompromised patients. The recent COVID-19 pandemic has underlined invasive fungal infections caused by *C.albicans*, particularly in immunocompromised individuals in hospital settings, as the major killer. Every year, more than 6,00,000 people are infected by *Candida sp.* Based on global data, the WHO Fungal Priority Pathogens List (2022) classifies *C. albicans* in the “Critical group” [1]. The pathogen can exist in yeast, pseudohyphae, and hyphae forms. The yeast-to-hyphae transition is mediated by many environmental cues involving different pathways. Transcription factors like Efg1p, Ace2p, Tup1p, and Cph2p regulate the transition from yeast to hyphae forms [2]. In addition, epigenetic modulators also modulate the transition. The SWI/SNF complex is one such modulator that controls the yeast-to-hyphae transition [3].

The SWI/SNF complex was purified as a ∼2 MDa complex from *S. cerevisiae*[4]. It comprises the catalytic subunit, Snf2, and at least 10-11 accessory subunits. The Snf2 protein belongs to the ATP-dependent chromatin remodeling protein family as it contains the seven conserved helicase motifs [5,6]. Homologs of the protein have been found in all organisms. The mammalian cells contain two homologs of *Sc*Snf2-BRG1 and BRM-that form distinct complexes [7,8]. The SWI/SNF complex repositions nucleosomes and, thus, regulates gene expression [9,10]. In *S. cerevisiae*, the SWI/SNF complex regulates ∼5% of the genes. In addition to its role in transcription, the complex also has been shown to play a role in the DNA damage response pathway [11–14]. The yeast SWI/SNF complex is required for initiating end resection during DNA double-strand break repair [15] while the mammalian SWI/SNF complex has been shown to recruit γH2AX to the double-strand breaks [13,14].

The SWI/SNF complex in *C. albicans*, like in *S. cerevisiae*, has 11 subunits [16,17]. The complex is required for hyphae formation [3], and is recruited to the promoters of the hyphae-specific genes via the transcription factor Efg1 [18]. The *C. glabrata Cgsnf2Δ* null mutants are deficient in biofilm formation [19]. In both *C. albicans* and *C. glabrata*, the SWI/SNF complex is required for virulence [3,19]. The complex is also a co-activator for the transcription factor, Mrr1, which regulates the expression of *MDR1*, a drug efflux pump encoding gene [20]. Studies have also shown that the complex is important for immune evasion in *C. glabrata* by regulating the transcriptional response within the macrophage [19]. However, the role of the complex in DNA damage response in *C. albicans* has not been studied.

Differences in the DNA damage response between *C. albicans* and *S. cerevisiae* have been shown by many studies. The DNA damage response pathway begins with the activation of the sensor kinases, Mec1 and Tel1. In *S. cerevisiae*, *MEC1* is essential. In contrast, *MEC1* in *C. albicans* is not essential, even though it mediates genome stability [21]. Tel1, in *S. cerevisiae*, is partially redundant with Mec1 and possesses a PI3Kc domain. The *C. albicans* Tel1 does not possess a PI3Kc domain, suggesting a potential functional diversity from *S. cerevisiae* [22]. The protein, though, has not been functionally characterised in *C. albicans*. Downstream of Tel1 and Mec1, Rad53, an effector kinase, is essential in *S. cerevisiae* but not in *C. albicans* [23]. However, the protein seems to have functional similarity as the loss of function of Rad53 leads to G2/M arrest both in *C. albicans* and *S. cerevisiae*.

The role of the SWI/SNF complex during DNA damage response and repair in *C. albicans* has not been fully explored. In this paper, we have studied the function of *Ca*Snf2 in modulating DNA damage response to genotoxic stressors. *Ca*Snf2 regulates the expression of DNA damage response genes as well as of *Ca*Fun30, an ATP-dependent chromatin remodeling protein, and CaRtt109, a histone acetyltransferase. The *Casnf2Δ* null mutant shows resistance to methyl methanesulfonate (MMS) and hydroxyurea (HU). MMS resistance is mediated by the upregulation of *CaRAD9,* an adaptor protein-encoding gene, by *Ca*Fun30 and *Ca*Rtt109. HU resistance appears to be driven by changes in the cell wall composition, leading to reduced uptake. Finally, we show that *Ca*Snf2 also regulates the expression of drug efflux pumps, thus, modulating the response of the pathogen to azoles.

## RESULTS

### *C. albicans SNF2* is required for growth

Studies have shown *S. cerevisiae SNF2* is not essential for viability and growth [24]. *Sc*Snf2p and *Ca*Snf2 share 51.9% identity. To understand the role of *C. albicans SNF2,* we first tagged one allele of the gene with a 6x His construct such that the expressed protein would have a 6x His tag that can be used for ChIP experiments. The strain was labeled as *CaSNF2His*. To study the role of *CaSNF2* in growth and viability, a null mutant (*Casnf2Δ*) was created in *CaSNF2His* background using the methodology delineated in Methods. Expression of protein was confirmed by western blot using anti-His antibody. As expected, the expression of the protein in the heterozygous strain (*Casnf2Hz*) was less as compared to the wild type (*CaSNF2His*) and the null strain (*Casnf2Δ*) did not show any Snf2 band (Fig. 1A (i) and (ii)).

**Figure 1.**
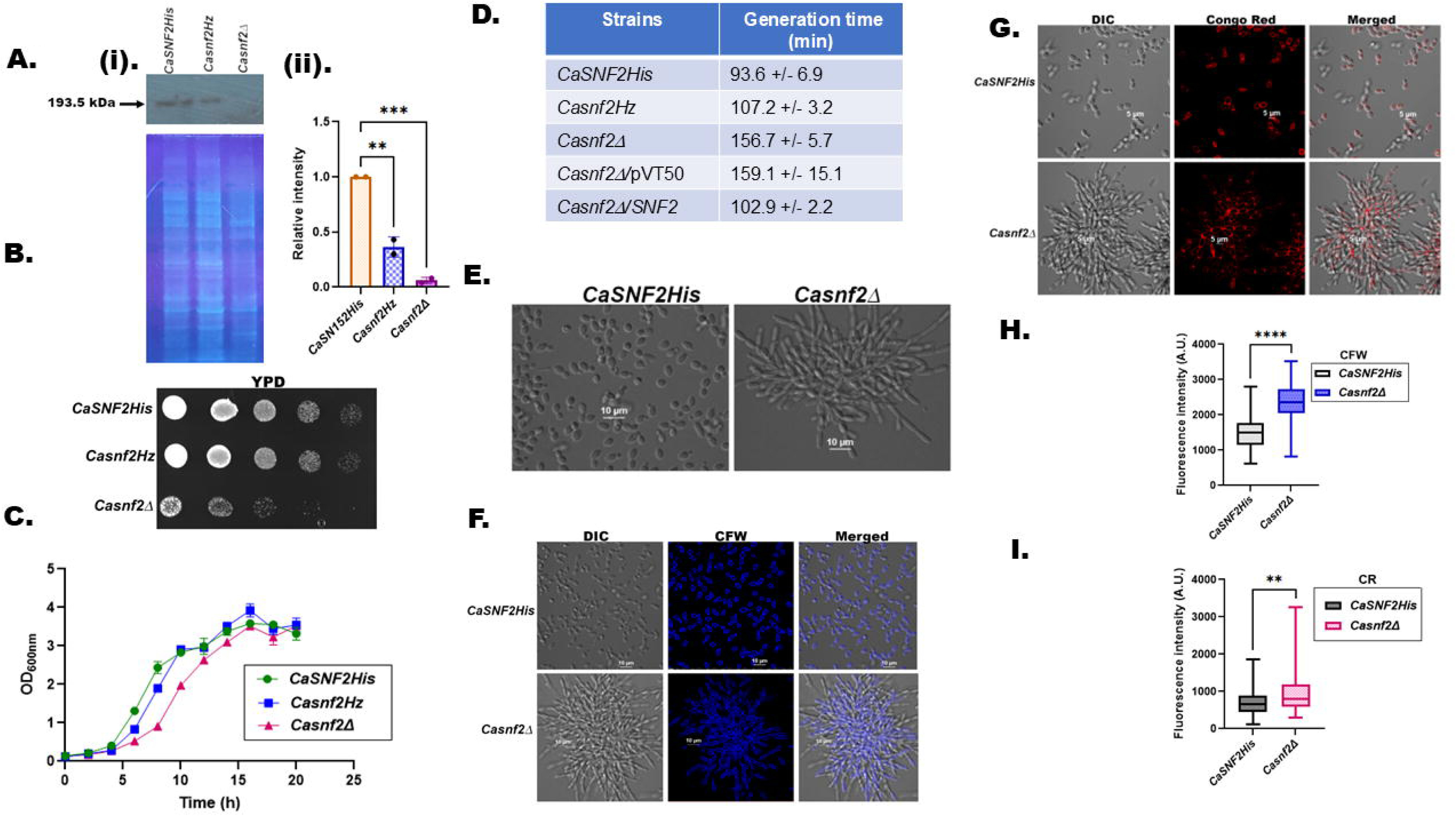
*C. albicans SNF2* is required for growth. (A). (i) Expression of CaSnf2 in wild type (*CaSNF2His*), heterozygous (*Casnf2Hz*), null (*Casnf2Δ*) was monitored using anti-His antibody. Stain-free technique (explained in materials and methods) was used to assess equal loading. (ii) Quantitation of the western blot. The data is presented as an average ± s.d. of two independent experiments. Normalization was done with respect to the gel visualized using stain-free technology as explained in materials and methods. ***indicates p = 0.0007 and ** indicates p = 0.024. (B). Plate assay monitoring growth of *CaSNF2His*, *Casnf2Hz,* and *Casnf2Δ* on YPD plate. Images were taken 24 h after incubation at 30°C. (C). Growth curve of *CaSNF2His*, *Casnf2Hz,* and *Casnf2Δ* in YPD media. (D). Generation times of *CaSNF2His*, *Casnf2Hz,* and *Casnf2Δ* on YPD plate in YPD media. (E). Confocal images of *CaSNF2His* and *Casnf2Δ*. (F). Calcofluor white (CFW) staining of *CaSNF2His* and *Casnf2Δ* cells. (G). Congo Red (CR) staining of *CaSNF2His* and *Casnf2Δ* cells. (H). Quantitation of the CFW fluorescence intensity. (I). Quantitation of the CR fluorescence intensity. Quantitation was done for >100 cells for each strain as explained in Materials and Methods. ***** indicates p< 0.0001. ** indicates p = 0.012.

The *Casnf2Δ* cells were viable, suggesting that, like *ScSNF2*, the gene is not essential. However, growth kinetics showed that the generation time for *Casnf2Δ* was ∼ 1.6-fold slower than the wild type (Fig. 1B-D). Analysis of the cells under the microscope showed that *Casnf2Δ* were clumped (Fig. 1D). Each cell was elongated as compared to the wild-type cell (Fig. 1E). The phenotype of the *Casnf2Δ* cells suggested that cell division was impaired. Therefore, chitin and β-glucan deposition were analyzed using calcofluor white (CFW) and Congo red (CR). Fluorescence intensity analysis showed that chitin and β-glucan levels were significantly higher (P<0.001) in the *Casnf2Δ* cells than in the wild-type cells (Fig. 1F-I).

To prove that the observed phenotypic changes were due to Snf2, a revertant strain (*Casnf2Δ/SNF2*) was created wherein *CaSNF2* was expressed in *Casnf2Δ* cells under the control of the constitutive promoter, p*ACT1*. As a control the vector, pVT50, was transfected into *Casnf2Δ* cells creating *Casnf2Δ/pVT50* control cell line. As expected, growth was restored in the *Casnf2Δ/SNF2* strain (Fig. 2A-F).

**Figure 2.**
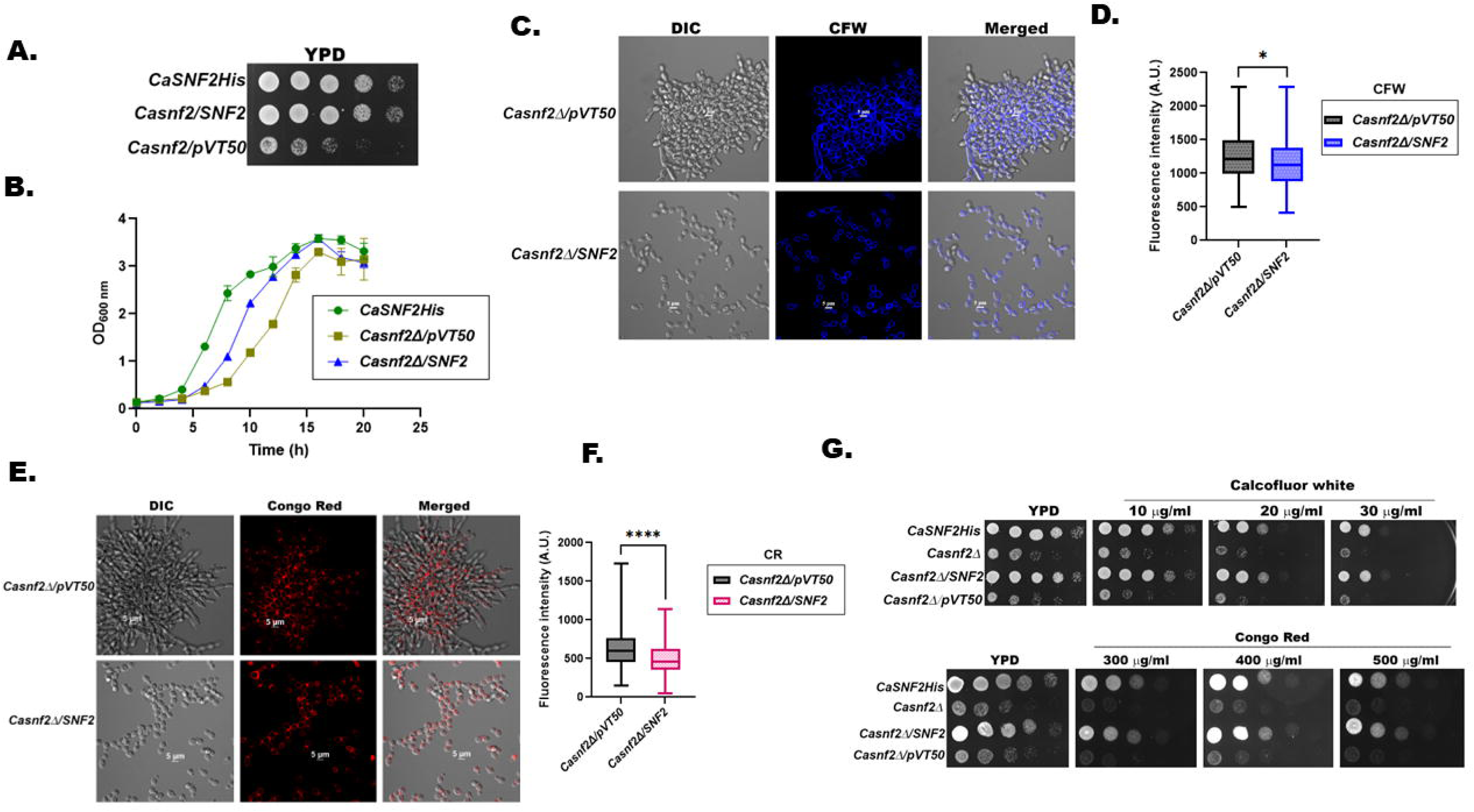
*CaSNF2* is required for cell wall integrity. (A). Plate assay monitoring growth of *CaSNF2His*, *Casnf2Δ/SNF2,* and *Casnf2Δ/pVT50* on YPD plate. Images were taken 24 h after incubation at 30°C. (B). Growth curve of *CaSNF2His*, *Casnf2Δ/SNF2,* and *Casnf2Δ/pVT50* in YPD media. (C). CFW staining of *Casnf2Δ/pVT50* and *Casnf2Δ/SNF2*cells. (D) Quantitation of the CFW fluorescence intensity. (E). CR staining of *Casnf2Δ/pVT50* and *Casnf2Δ/SNF2*cell. (F). Quantitation of the CR fluorescence intensity. (G). Plate assays monitoring the growth of *CaSNF2His*, *Casnf2Δ Casnf2Δ/SNF2,* and *Casnf2Δ/pVT50* on CFW and CR plates. Images were taken 48 h after incubation at 30°C. Quantitation of CFW and CR was done for >100 cells for each strain as explained in Materials and Methods. * Indicates p = 0.04 and ***** indicates p< 0.0001.

### CaSNF2 is required for the cell wall integrity

As *Casnf2Δ* mutant cells showed increased chitin and β-glucan levels, the response of these cells to CFW and CR in a plate assay was analyzed. The *Casnf2Δ* mutant cells were more sensitive to these anionic dyes than the wild-type cells indicating *CaSNF2* is required for cell wall formation (Fig. 2G).

### *Ca*Snf2 regulates the expression of *DBF2, ACE2*, and *MOB1*

Ace2 is a zinc-finger transcription factor that regulates the expression of genes related to cell wall separation and integrity. Dbf2, in *S. cerevisiae*, is required for telophase/G1 transition while Mob1 interacts with Dbf2 to direct the late mitotic events in the cell [25,26]. *Casnf2Δ s*howed cell division defects, so we hypothesized that the protein might co-regulate *CaACE2*, *CaDBF2*, and *CaMOB1* expression. The hypothesis was confirmed as qPCR analysis showed that the expression *CaACE2*, *CaDBF2*, and *CaMOB1* were downregulated in *Casnf2Δ* cells compared to the wild type (Fig. 3A). Further, the expression was restored in *Casnf2Δ/SNF2* cells (Fig. 3B). ChIP experiments confirmed occupancy of *Ca*Snf2 and RNAPII on the promoter of *CaACE2* (Fig. 3C).

**Figure 3.**
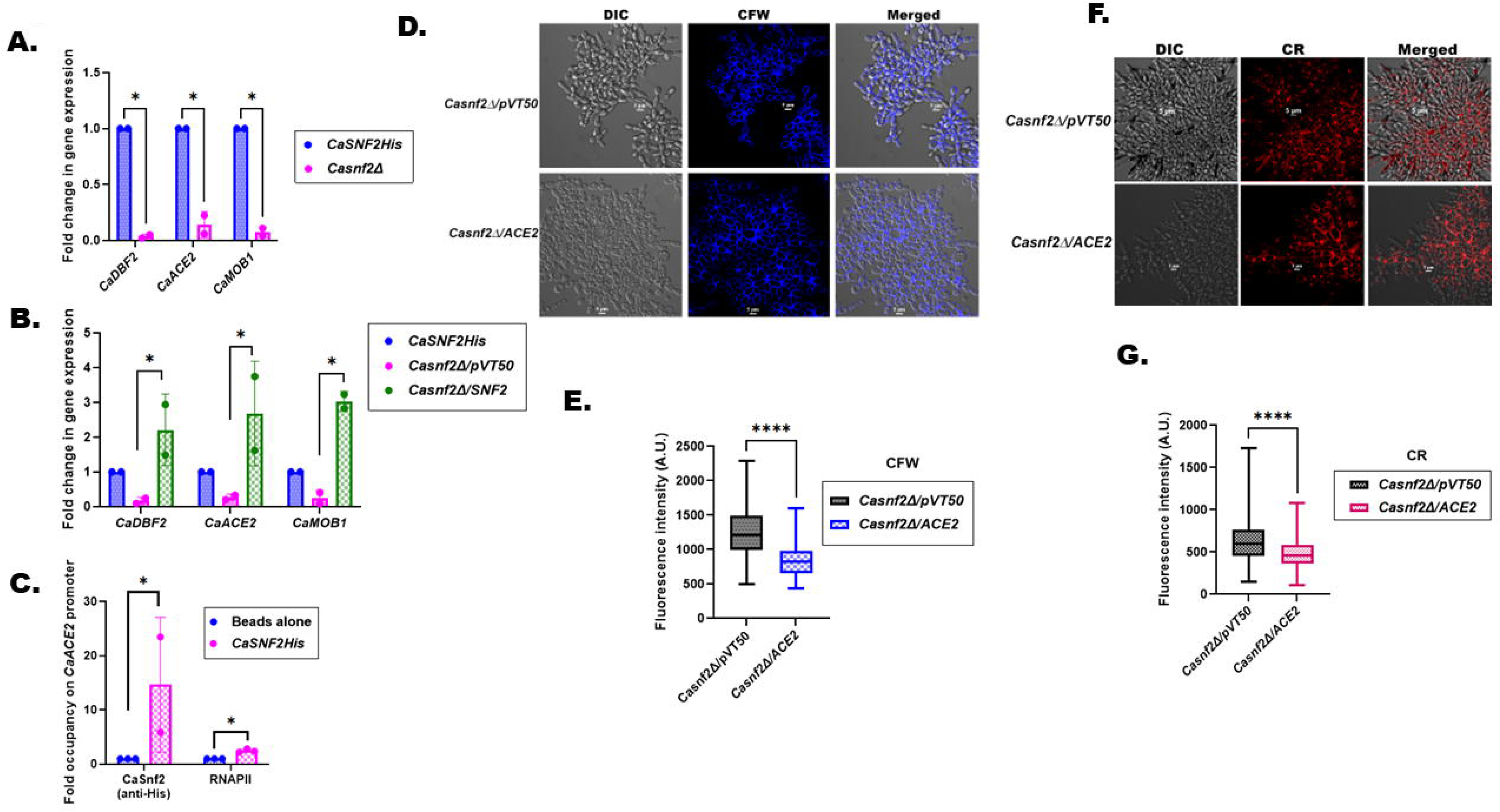
CaSnf2 regulates the expression of *CaDBF2, CaACE2*, and *CaMOB1*. (A). Expression of *CaDBF2, CaACE2*, and *CaMOB1* in *CaSNF2His* and *Casnf2Δ* cells. (B). Expression of *CaDBF2, CaACE2*, and *CaMOB1* in *CaSNF2His*, *Casnf2Δ*/*pVT50*, and *Casnf2Δ/SNF2* cells. (C). Occupancy of CaSnf2 and RNAPII on *CaACE2* promoter. The occupancy of CaSnf2 was probed using an anti-His antibody. (D). CFW staining of *Casnf2Δ/pVT50* and *Casnf2Δ/ACE2*cells. (E) Quantitation of the CFW fluorescence intensity. (F). CR staining of *Casnf2Δ/pVT50* and *Casnf2Δ/ACE2*cells. (G). Quantitation of the CR fluorescence intensity. The qPCR and ChIP data are the average ± sem of two independent experiments. The star indicates p< 0.05. Quantitation of CFW and CR was done for >100 cells for each strain as explained in Materials and Methods. ***** indicates p< 0.0001.

Next, we asked whether overexpression of *CaACE2* in *Casnf2Δ* would rescue the cell division and cell wall defects. Confocal analysis showed that *Casnf2Δ/CaACE2* cells had a flatter appearance with a decrease in chitin and β-glucan deposition. However, the cell division defects persisted as can be observed by cell clumping (Fig. 3D-G).

Thus, overexpression of *CaACE2* does not rescue the cell division defects observed in *Casnf2Δ* cells. This is not surprising as Ace2 mediated transcription might need Snf2 (or SWI/SNF complex) as a co-regulator.

### *CaSnf2* regulates the expression of DNA damage response genes

Mammalian BRG1 has been shown to regulate the expression of *ATM* and *ATR*, the genes encoding the sensor kinases [27–30]. Therefore, we asked whether *C. albicans* Snf2 also regulates the expression of DNA damage response genes. The expressions of *CaTEL1* and *CaMEC1*, *CaMRC1*, *CaRAD9*, *CaFUN30*, *CaRTT109*, and *CaRAD5* were analyzed. CaTel1 and CaMec1 are the homologs of ATM and ATR in fungi. Mrc1[31,32] and Rad9[33], two adaptor proteins phosphorylated by Tel1 and Mec1, play a role in recruiting repair proteins to the DNA damage site. Previous studies have shown that Fun30, an ATP-dependent chromatin remodeling protein, and Rtt109, a fungal-specific histone acetyltransferase, regulate the expression of *CaSNF2*[34]. To understand whether a feedback loop exists, the expressions of *CaFUN30* and *CaRTT109* were quantitated. Finally, the expression of *CaRAD5* that is required for post-replication repair [35] was also analyzed.

qPCR analysis showed that the expression of *CaSNF2, CaFUN30, CaRTT109, CaTEL1, CaMEC1, CaMRC1, CaRAD9* and *CaRAD5* were downregulated in *Casnf2Δ* cells as compared to the wild-type cells (Fig. 4A). To determine the specificity of the qPCR analysis, the expression levels of *CaEFB1* and *CaPMA1* were studied. *CaEFB1* codes for the elongation factor EF-1β[36] while *CaPMA1*[37] codes for the proton pump, Pma1p, involved in growth and pH homeostasis. The expression of both these genes was found to be unchanged in *Casnf2Δ* suggesting *Ca*snf2 regulates the expression of only a subset of genes.

**Figure 4.**
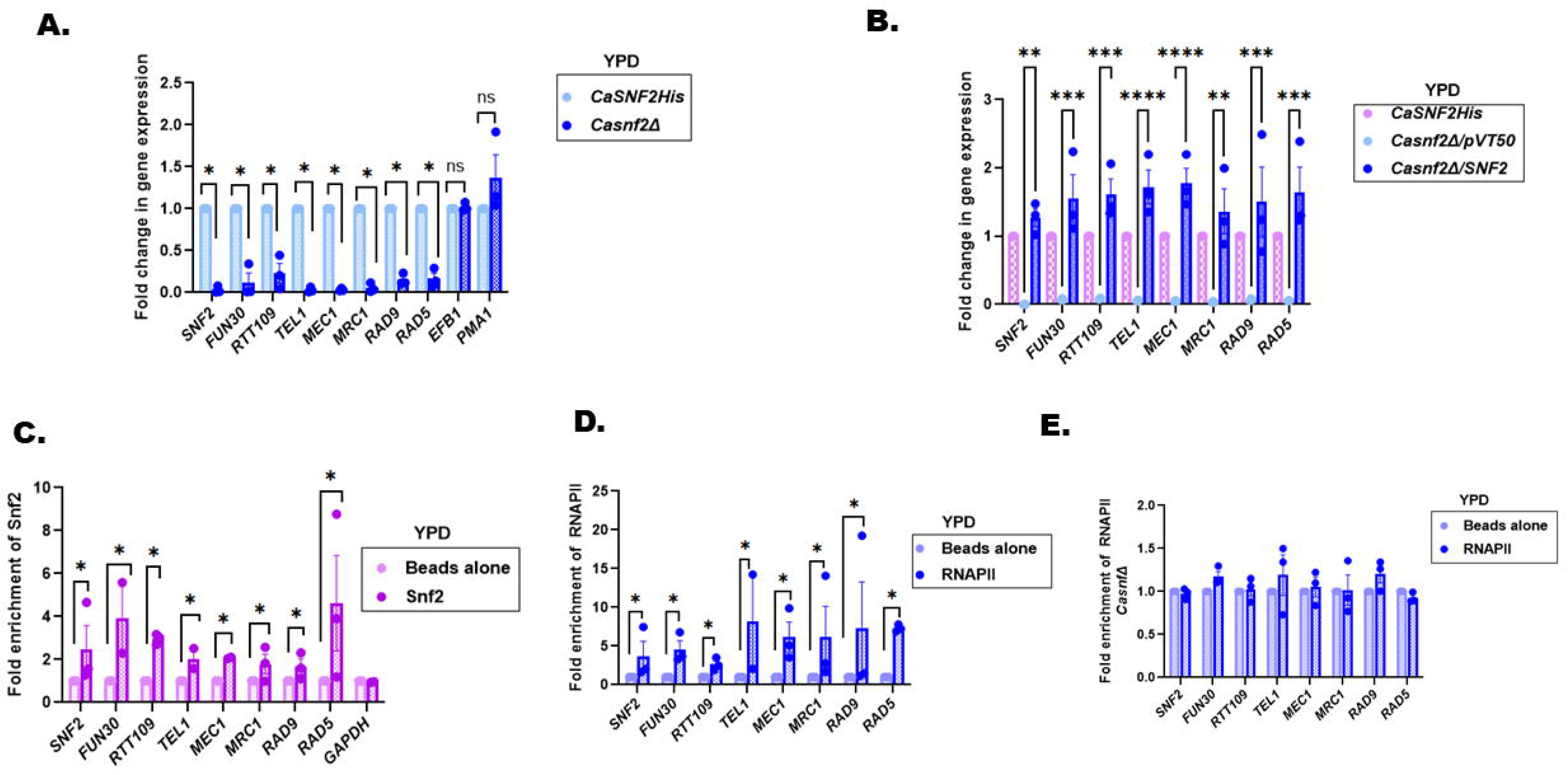
*CaSnf2* regulates the expression of DNA damage response genes. (A). Expression of *CaSNF2, CaFUN30, CaRTT109, CaTEL1, CaMEC1, CaMRC1, CaRAD9, CaRAD5, CaEFB1,* and *CaPMA1* were analyzed in *CaSNF2His* and *Casnf2Δ* cells. (B). Expression of *CaSNF2, CaFUN30, CaRTT109, CaTEL1, CaMEC1, CaMRC1, CaRAD9,* and *CaRAD5* were analyzed in *CaSNF2His*, *Casnf2Δ/pVT50, Casnf2Δ/SNF2* cells. (C). Occupancy of *Ca*Snf2 on *CaSNF2, CaFUN30, CaRTT109, CaTEL1, CaMEC1, CaMRC1, CaRAD9, CaRAD5,* and *CaGAPDH* promoters were analyzed using anti-His antibody in *CaSNF2His* cells. (D). Occupancy of RNAPII on *CaSNF2, CaFUN30, CaRTT109, CaTEL1, CaMEC1, CaMRC1, CaRAD9,* and *CaRAD5* was analyzed in *CaSNF2His* cells. (E). Occupancy of RNAPII on *CaSNF2, CaFUN30, CaRTT109, CaTEL1, CaMEC1, CaMRC1, CaRAD9,* and *CaRAD5* was analyzed in *Casnf2Δ* cells. The qPCR and ChIP data are the average ± sem of three independent experiments. * indicates p<0.05; ** indicates p = 0.003; *** indicates p = 0.0004 and **** indicates p<0.0001.

To further confirm the specificity of regulation, the expression of *CaSNF2, CaFUN30, CaRTT109, CaTEL1, CaMEC1, CaMRC1, CaRAD9,* and *CaRAD5* were analyzed in *Casnf2Δ/SNF2*. As compared to the control, *Casnf2Δ/pVT50*, the expression of all these genes was upregulated in *Casnf2Δ/SNF2* confirming that *Ca*Snf2 either directly or indirectly regulates the expression of the DNA damage response genes (Fig. 4B).

ChIP experiments were performed using anti-His antibody to study the occupancy of *Ca*Snf2 on the promoters of *CaSNF2, CaFUN30, CaRTT109, CaTEL1, CaMEC1, CaMRC1, CaRAD9,* and *CaRAD5*. Using the same method, the occupancy of RNAPII on these promoters was also investigated. *Ca*Snf2 and RNAPII were present on all the promoters, indicating that the ATP-dependent chromatin remodeler directly regulates the expression of these genes (Fig. 4C and D). Further, RNAPII occupancy on these genes unchanged in *Casnf2Δ/SNF2* cells as compared to *Casnf2Δ/pVT50* cells, correlating with downregulated expression in the mutant cells (Fig. 4E). To determine the specificity, occupancy of *Ca*Snf2 on the *CaGAPDH* promoter was probed (Fig. 4C). The protein was not found to be enriched on this promoter.

From these experiments, it was concluded that *Ca*Snf2 regulates the expression of DNA damage response genes and is needed for RNAPII recruitment to the promoter of these genes.

### *CaSNF2* is not required for CPT-mediated DNA damage response

As *Ca*Snf2 regulates the expression of DNA damage response genes, the role of the protein in DNA damage response was next analyzed. Plate assays showed that *Casnf2Δ* cells are phenotypically like the wild-type cells (Supplementary Fig. 1). This result was surprising as both BRG1 in humans and Snf2 in *S. cerevisiae* are required for DNA double-strand break repair [38,39] . Gene expression analysis showed the expression of *CaFUN30, CaRTT109, CaTEL1, CaMEC1, CaMRC1, CaRAD9,* and *CaRAD5* was upregulated in the presence of CPT in mutant cells, though the expression was still only 50% compared to the wild-type cells (Supplementary Fig. 1). Further, the genes involved in NHEJ pathway (*YKU80*, *CAS1*, and *LIG4*) were also only 50% downregulated as compared to the wild type cells (Supplementary Fig. 1).

From these results, it was concluded that *CaSNF2* expression is not required to repair CPT-induced DNA damage probably because the expression of the DNA damage response genes was sufficient to drive the repair process.

### Resistance to MMS in *Casnf2Δ* mutant is mediated by *CaRAD9* upregulation by *CaFun30* and *CaRtt109*

Next, the role of *C*aSnf2 in mediating response to mismatches was studied using MMS. Plate assays and MIC80 showed that *Casnf2Δ* cells are resistant to MMS compared to the control cells (Fig. 5A-C). The resistant phenotype was reversed in *Casnf2Δ/SNF2* (Fig. 5A-C). These results were contrary to *S. cerevisiae* where *snf2Δ* cells do not show any phenotypic response to MMS compared to the wild type [24].

**Figure 5.**
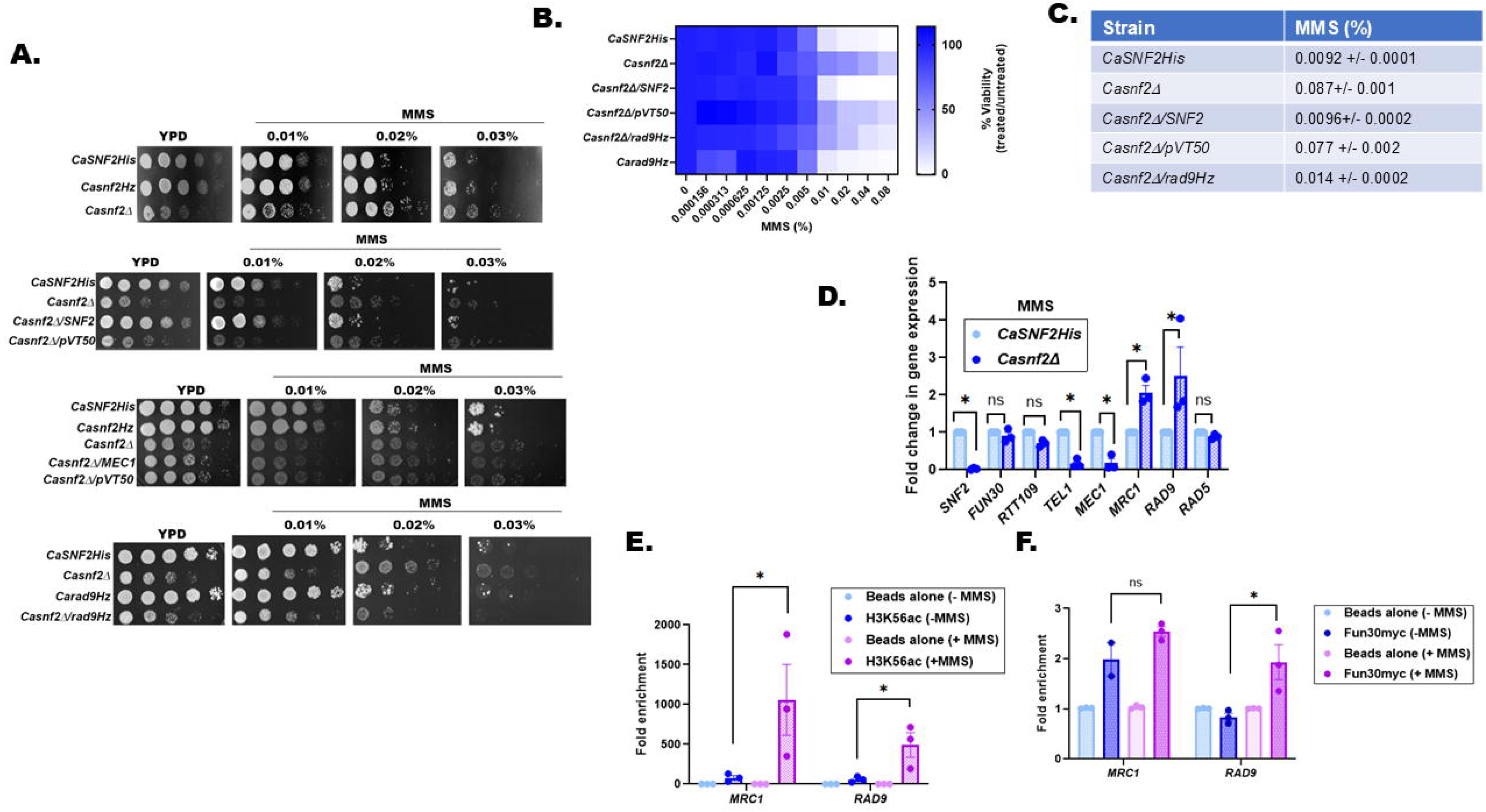
Resistance to MMS in *Casnf2Δ* mutant is mediated by *CaRAD9* upregulation by *CaFun30* and *CaRtt109*. (A). Plate assays showing the response of *CaSNF2His* and mutant strains to MMS. (B). Heat map showing the response of the wild type and mutant strains to increasing concentration of MMS. (C). MIC80 of wild type and mutant strains of *C. albicans*. (D). Expression of *CaSNF2, CaFUN30, CaRTT109, CaTEL1, CaMEC1, CaMRC1, CaRAD9,* and *CaRAD5* were analyzed in the presence of MMS in *CaSNF2His* and *Casnf2Δ* cells. (E). Occupancy of H3K56ac on *MRC1* and *RAD9* promoter in *Casnf2Δ* cells in the absence and presence of MMS. (F). Occupancy of Fun30 on *MRC1* and *RAD9* promoter in *Casnf2Δ* cells in the absence and presence of MMS. One copy of *CaFUN30* was tagged with myc in *Casnf2Δ* cells such that the expressed protein was myc tagged. Anti-myc antibody was used for the ChIP experiments. The qPCR data is the average ± sem of three independent experiments. The ChIP data is the average ± sem of two independent experiments. The star indicates p< 0.05.

Gene expression analysis revealed that expression of *CaMEC1* was downregulated while *CaMRC1* and *CaRAD9* expression was upregulated in *Casnf2Δ* (Fig. 5D). This was reminiscent of the phenotype and gene expression pattern observed in *CaFUN30Hz/CaRTT109Hz* cells, which also showed resistance to MMS and similar expression pattern of these three genes [34].

As *CaMEC1* is the sensor kinase that phosphorylates both Mrc1 and Rad9, we first investigated whether overexpression of this kinase gene would revert the phenotype. However, *Casnf2Δ/MEC1* was not able to rescue the resistant phenotype (Fig. 5A). Next, the role of *Ca*Rad9 was investigated by deleting this gene in *Casnf2Δ* background creating *Casnf2Δ/rad9Hz* mutants. Plate assays showed that *Casnf2Δ/rad9Hz* was able to revert the resistance phenotype, suggesting *CaRAD9* does play an important role in mediating response to MMS (Fig. 5A-C).

ChIP experiments showed that the occupancy of both *Ca*Fun30 (probed using anti-Myc antibody) and *Ca*Rtt109 (probed using anti-H3K56ac antibody) increased on *CaRAD9* promoter on MMS treatment (Fig. 5E-F).

From these experiments, it was concluded that *Ca*Fun30 and *Ca*Rtt109 regulate the response to MMS by modulating the expression of *CaRAD9*.

### Resistance to HU in *Casnf2Δ* mutant is due to cell wall perturbations

Finally, the response of *Casnf2Δ* to HU was studied. Plate assays, broth dilution assays, and MIC80 showed that *Casnf2Δ* cells are resistant to HU compared to wild-type cells (Fig. 6A-C). The resistant phenotype was reversed in *Casnf2Δ/SNF2* (Fig. 6A-C). Again, this result was contrary to *S. cerevisiae* where *snf2Δ* mutants were found sensitive to HU [24,40].

**Figure 6.**
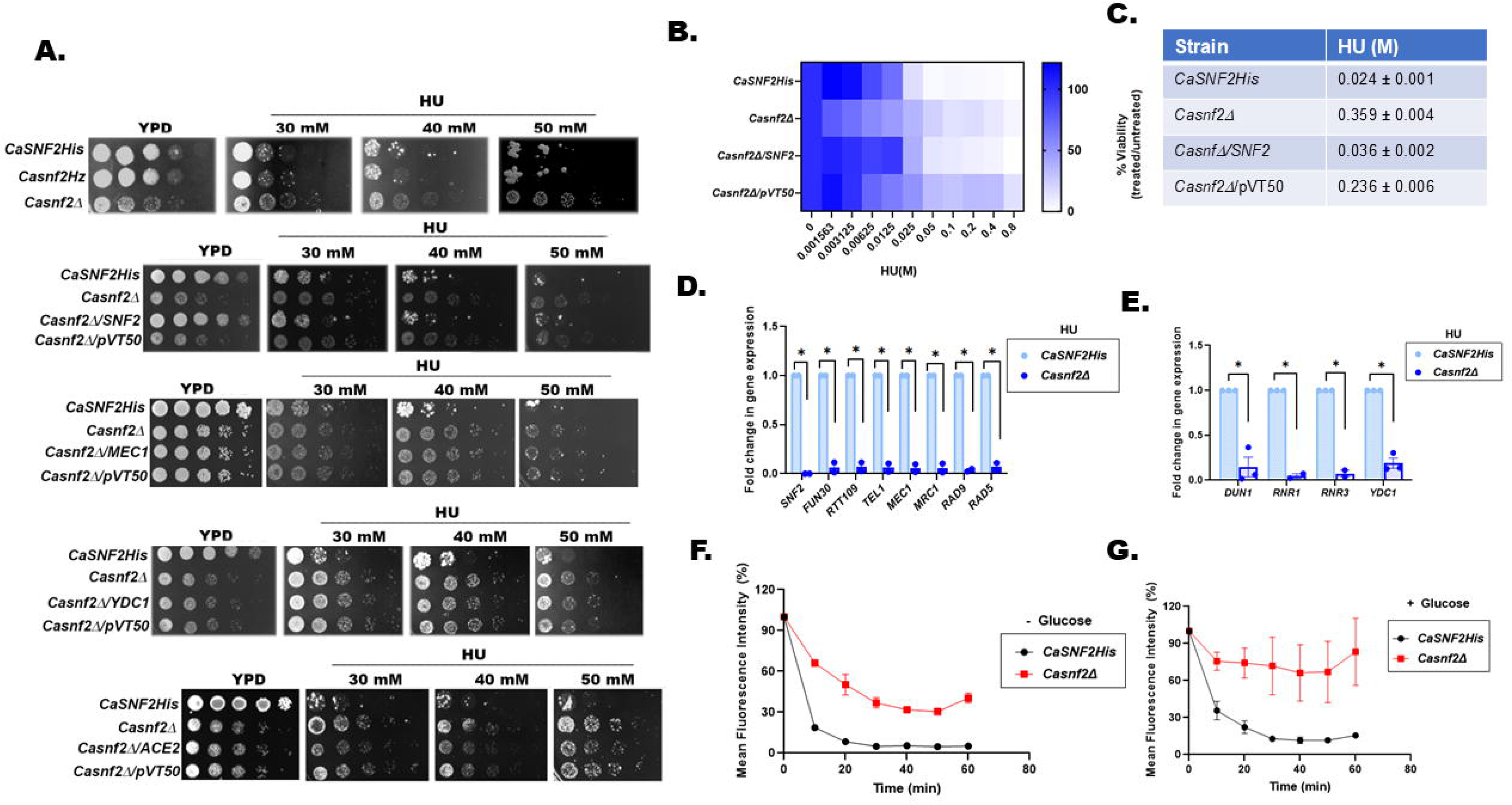
Resistance to HU in *Casnf2Δ* mutant is due to cell wall perturbations. (A). Plate assays showing the response of *CaSNF2His* and mutant strains to HU. (B). Heat map showing the response of the wild type and mutant strains to increasing concentration of HU. (C). MIC80 of wild-type and mutant strains of *C. albicans*. (D). Expression of *CaSNF2, CaFUN30, CaRTT109, CaTEL1, CaMEC1, CaMRC1, CaRAD9,* and *CaRAD5* was analyzed in the presence of HU in *CaSNF2His* and *Casnf2Δ* cells. The experiment is presented as average ± sem of two independent experiments. (E). Expression of *CaDUN1*, *CaRNR1*, *CaRNR3*, and *CaYDC1* was analyzed in the presence of HU in *CaSNF2His* and *Casnf2Δ* cells. The qPCR data is the average ± sem of three independent experiments. (F). Rhodamine 6G uptake was measured in the absence of glucose as a function of time in *CaSNF2His* and *Casnf2Δ* cells. (G). Rhodamine 6G uptake was measured in the presence of glucose as a function of time in *CaSNF2His* and *Casnf2Δ* cells. The Rhodamine 6G uptake assay is the average ± sd of two independent experiments. The star indicates p< 0.05.

Gene expression analysis showed that *CaSNF2, CaFUN30, CaRTT109, CaTEL1, CaMEC1, CaMRC1, CaRAD9,* and *CaRAD5* was downregulated in *Casnf2Δ* cells in the presence of HU (Fig. 6D). HU inhibits ribonucleotide reductase (RNR). Therefore, we hypothesized that possibly RNR genes are upregulated in *Casnf2Δ* mutant providing resistance to HU. We also monitored the expression of *CaDUN1* which encodes Dun1, a positive regulator of RNR. However, *CaDUN1, CaRNR1,* and *CaRNR3* were downregulated in the mutant cells in the presence of the genotoxic stressor (Fig. 6E). In *S. cerevisiae*, ceramides have been shown to play a role in HU resistance [41]. Therefore, we analyzed the expression of *CaYDC1*, which is the homolog of *ScYDC1*, and found it to be downregulated in *Casnf2Δ* cells in the presence of HU (Fig. 6E).

Based on the qPCR data, we hypothesized that HU resistance in the mutant cells was driven either by *CaMEC1* or *CaYDC1* downregulation. However, neither *Casnf2Δ/MEC1* nor *Casnf2Δ/YDC1* rescued the resistant phenotype (Fig. 6A).

HU treatment has been shown to induce polarized growth in *C. albicans* [42] Therefore, the cells were stained with CFW and examined using a confocal microscope. The wild-type and *Casnf2Δ* mutants showed polarized growth; however, the length of the mutant cells was less (P value = 0.0001) compared to the wild-type (Supplementary Fig.2).

Based on this result, we hypothesized that the HU resistance could be due to the accumulation of chitin and β-glucan on the cell wall of the *Casnf2Δ* mutant. Therefore, we examined the response of *Casnf2Δ/ACE2* (where both chitin and β-glucan levels were reduced) to HU and found that overexpression of *CaACE2* in the *Casnf2Δ* mutant partially restored the phenotype (Fig. 6A).

Finally, the Rhodamine-6G uptake assay performed both in the absence and presence of glucose showed that the uptake of the dye was impaired in the *Casnf2Δ* mutant compared to the wild-type cells (Fig. 6F and G).

From these results, we hypothesize that *Casnf2Δ* mutants are resistant to HU due to altered chitin and β-glucan levels on the cell wall, leading to decreased uptake of the genotoxic stressor.

### *Casnf2Δ* mutants are sensitive to azoles

Azoles are the primary drugs for Candidiasis treatment. We hypothesized that *Casnf2Δ* mutants should be resistant to azoles as the uptake of Rhodamine-6G was reduced in these cells compared to the wild-type. Surprisingly, plate assays and MIC80 showed that the mutant cells were sensitive to azoles (Fig. 7A-C).

**Figure 7.**
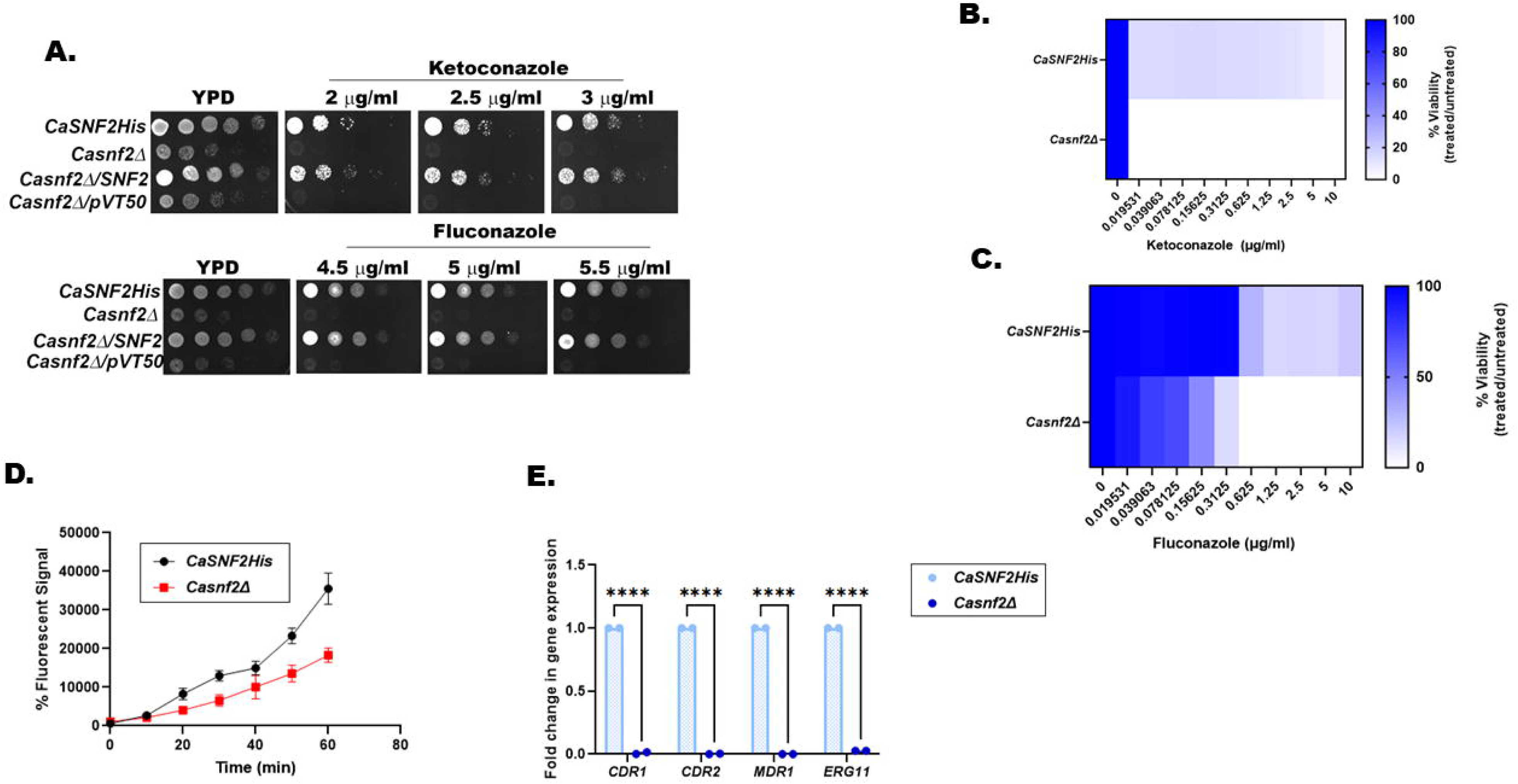
*Casnf2Δ* mutants are sensitive to azoles. (A). Plate assays showing the response of *Casnf2Δ* to ketoconazole and fluconazole. (B). Heat map showing the response of *Casnf2Δ* cells to increasing concentration of ketoconazole. (C). Heat map showing the response of *Casnf2Δ* cells to increasing concentration of fluconazole. (D). Rhodamine 6-G efflux was measured as a function of time in *CaSNF2His* and *Casnf2Δ* cells. The efflux assay is the average ± sd of two independent experiments. (E). Expression of *CaCDR1, CaCDR2, CaMDR1,* and *CaERG11* was analyzed in *CaSNF2His* and *Casnf2Δ* cells. The qPCR data is the average ± sem of two independent experiments. The star indicates p< 0.05.

The Rhodamine-6G assay showed reduced efflux of the dye (Fig. 7D). Further, gene expression analysis showed that the expressions of *CaCDR1*, *CaCDR2*, and *CaMDR1* were downregulated in *Casnf2Δ* cells compared to the wild-type cells. The expression of *ERG11*was also downregulated (7E).

From these results, it was concluded that *Casnf2Δ* cells are sensitive to azoles due to the downregulation of the efflux pumps.

## DISCUSSION

The ATP-dependent chromatin remodeling factor Snf2 mediates a diverse range of activities in the cell. Using the energy released from ATP hydrolysis, the protein repositions nucleosomes and thus modulates the expression of ∼5% of genes in *S. cerevisiae*. It also regulates DNA damage response and replication [11,12]. In *Candida sp.* the protein has been shown to regulate filamentation and virulence. In this paper, we have explored the role of *C*aSnf2 in DNA damage response in *C. albicans*.

*C. albicans* Snf2 shows functional divergence from *S. cerevisiae* Snf2 despite sharing approximately 52% homology. *CaSNF2* is required for cell division as the null mutant showed increased duplication time. The protein, like *Ca*Snf6, a component of the SWI/SNF complex, regulates expression of *Ca*Ace2 [16]. Further, chitin and β-glucan levels were upregulated in null mutants as compared to the wild type cells. *Ca*Snf2 appears to regulate deposition of chitin and β-glucan via *Ca*Ace2, a transcription factor, as overexpression of *CaACE2* in *Casnf2Δ* mutant restored chitin and β-glucan levels. However, the cell division defects persisted indicating that *Ca*Snf2 is required for the regulation of the cell division genes possibly by co-regulating with *Ca*Ace2. In *S. cerevisiae*, Snf2 regulates the cell wall integrity by regulating the MAPK pathway [43]. The role of *CaS*nf2 in modulating the MAPK pathway needs to be investigated.

Gene expression analysis showed that *Ca*Snf2 also regulates the expression of DNA damage repair genes: *CaTEL1*, *CaMEC1*, *CaRAD9* and *CaMRC1*. *S. cerevisiae* Snf2 is known to regulate the activity of *Sc*Mec1[40]. In *C. albicans*, Snf2 also appears to regulate its expression at transcriptional level. In addition, the expression of two ATP-dependent chromatin remodeling proteins-*CaRAD5* and *CaFUN30*-as well as of *CaRTT109*, a fungal-specific histone acetyltransferase, also appear to be regulated by *Ca*Snf2.

Based on the qPCR and ChIP experiments, the *Casnf2Δ* mutants were expected to be sensitive to genotoxic stressors. However, spot assays demonstrated that *Casnf2Δ* mutants are resistant to both MMS and HU as compared to the wild type cells. These results showed the difference between *Casnf2Δ* and *Scsnf2Δ* mutants, highlighting the dissimilarity in the function of this protein in the two organisms.

MMS activates mismatch repair pathway. In *S. cerevisiae*, Snf2 does not appear to be in the pathway conferring resistance to MMS and therefore, *Scsnf2Δ* mutants do not show any phenotypic response to this DNA damaging agent [24]. In *C.albicans*, the deletion of *SNF2* causes MMS resistance. Mechanistically, this resistance appears to be due to two factors. This first reason is the upregulation of *CaRAD9* by *Ca*Fun30 and *Ca*Rtt109 in *Casnf2Δ* cells. Deletion of one copy of *CaRAD9* in *Casnf2Δ* rendered the cells sensitive to MMS like the wild type cells. The second factor contributing to MMS resistance could be the increased level of chitin and β-glucan levels that lead to reduced uptake. Rhodamine-6-G uptake assay confirmed that the uptake of the fluorescent dye was less in the mutant cells as compared to the wild type cells both in the absence and the presence of glucose. The cell wall defect due to altered chitin and β-glucan levels is possibly also the cause of HU resistance in *Casnf2*Δ cells. In case of *S. cerevisiae*, *Scsnf2Δ* mutants are sensitive to HU [12,24]. The differential response of *Casnf2Δ* and *Scsnf2Δ* mutants to genotoxic stressors might lie in the fact that the pathogen can reside within the macrophages where it needs to counter the DNA damage due to oxidative stress.

Finally, *Casnf2Δ* mutants are sensitive to azoles due to downregulation of the multi-drug efllux pumps, making this protein an attractive drug target. Transcriptional regulation of the multi-drug resistant pumps by Snf2 appears to be a conserved feature as BRG1 too has been shown to regulate the expression of drug efflux pumps in breast cancer cell lines [44]. The future drug discovery program can either target the ATPase domain of *Ca*Snf2 or target the interaction between Snf2 and other components of the SWI/SNF complex.

## EXPERIMENTAL PROCEDURE

### Chemicals

All chemicals used in this study were of analytical grade and were purchased either from Thermo Fisher Scientific (USA), Merck (India), HiMedia (India), SRL (India), or Sigma-Aldrich (USA). The media for growing cells was purchased from HiMedia (India). Restriction enzymes and T4 DNA ligase were purchased from Thermo Fisher Scientific (USA), Merck (USA), and New England Biolabs (USA). The TA cloning kit was purchased from MBI Fermentas (USA). The gel extraction kit was purchased from Qiagen (Germany). SYBR Green was purchased from Kapa Biosystems (Switzerland). Bradford dye for protein estimation and Protein G agarose was purchased from Sigma-Aldrich (USA). Protein and DNA molecular weight markers were purchased from MBI Fermentas (USA). Primers were synthesized by GCC Biotech (India).

### Vectors and Strains

*C. albicans* strain SN152 strain was a kind gift from K. Natarajan, School of Life Sciences, JNU. The vector pVT50 used for cloning was a gift from Prof. Alok Mondal, School of Life Sciences, JNU, India. The vectors, pSN40 and pSN69, used for gene tagging were kind gift from Prof. K. Natarajan, School of Life Sciences, JNU, India. The primers used to construct mutants are listed in Supplementary Table 1.

### Cell culture

Yeast Extract-Peptone-Dextrose (YPD) media or synthetic dextrose (SD) minimal media supplemented with 85.6 g/ml histidine, or arginine or leucine was used to grow *C. albicans*. The list of strains used in this study is provided in Supplementary Table 2.

### Antibodies

Anti-H3K56ac (Cat# 76307) and anti-RNAPII (Cat# Ab5408) antibodies were purchased from Abcam (Cambridge, UK). Anti-c-Myc (Cat# ITG0001) and anti-His (Cat# M1001020) antibodies were purchased from Immunotag (St. Louis, MO, USA).

### Epitope-Tagging of *CaSNF2*

pSN40 vector w used to add 6x His and c-myc tags to *CaSNF2* and *CaFUN30* respectively. Briefly, the fusion PCR method was used to create an amplicon containing 5’ of the targeted gene, a selection marker, and 3’ of the targeted gene of the targeted gene [45]. The amplified fragment was transformed into the SN152 strain and transformants were selected using either His^-^ or Arg^-^ selection media. The 6X His tagged strain was termed *CaSNF2His* and the c-myc tagged strain was named *CaFUN30myc* (Supplementary Table 2).

### Generation of mutant strains

PCR-based gene deletion strategy was used for making the mutants [45]. Heterozygous mutants of *CaSNF2, CaMRC1* and *CaRAD9* were made by replacing one of the alleles of the respective genes with the selection marker *HIS1*. The *Casnf2* null mutant (*Casnf2Δ*) was created by replacing the second allele of *CaSNF2* with an *ARG4* selection marker. The *Casnf2*Δ*/CaRAD9Hz* was made by disrupting one of the alleles of *RAD9* with a *LEU2* selection marker in the *Casnf2*Δ background.

All the mutants were confirmed by PCR using gene-specific flanking primers (Supplementary Table 1).

### Generation of overexpression and revertant strains

The pVT50 vector was used to create the revertant and overexpression strains. The genes were amplified using PCR (Supplementary Table 3) and cloned between XhoI and NheI restriction sites, and the constructs were linearized using StuI. *Casnf2Δ* null strain was transformed with the linearized plasmid, and the transformants were selected on YPD plates containing 100 μg/ml nourseothricin. The successful transformants were screened by qPCR.

### SDS-PAGE

A single colony was inoculated in YPD media and grown for 16 h. 1 ml of primary culture was inoculated in 100 fresh YPD and the secondary culture was grown to 1 OD600 nm. The cells were harvested by centrifuging at 6000 rpm for 5 min and pellet was resuspended in 400 μl lysis buffer (40 mM Tris.Cl pH 7.0; 350 mM NaCl; 0.1% Tween-20; 10% glycerol; 1 mM PMSF; 50 mM NaF; 1X protease inhibitor complex) and incubated on ice for 30 min. Subsequently, glass beads were added and wild type and *Casnf2Hz* cells were lysed by vortexing (30 sec ON; 30 sec OFF; 20 cycles). The null mutant (*Casnf2Δ*) were lysed by vortexing for 30 cycles (30 sec ON; 30 sec OFF). The lysed samples were clarified by centrifuging at 13000 rpm for 15 min. The supernatant was again clarified by centrifuging at 13000 rpm for 10 min. Protein concentration was estimated using Bradford reagent. For western blotting, 300 μg protein was loaded on 10% SDS-PAGE containing 0.05% of 2,2,2-Tricholoethanol (Tokyo Chemical Industry Co, Ltd (Japan)) in the resolving gel. 2,2,2-Trichloroethanol covalently binds to tryptophan residues that fluoresces on exposure to UV light and was used to monitor equal loading (Stain-free technique) [46].

### Western blotting

After ensuring equal loading, the proteins were transferred to PVDF membrane at 850V for 210 min. The blots were blocked for 45 min using blocking buffer containing 3% skim milk in TBST. After blocking, the membranes were incubated in primary antibody for 18 hr at 4°C. Subsequently, the membranes were incubated at room temperature for 3 h. After incubation, the blots were washed twice with TBST for 5 min each. The membranes were incubated in secondary antibody (1: 3000) for 2 hr at room temperature. Subsequently, the blots were washed twice with TBST for 5 min and developed using ECL reagent

### Quantitative real-time RT-PCR (qPCR)

Cells from the log phase of *Candida albicans* culture were centrifuged, washed with DEPC-treated water, and RNA was extracted using 500 µl of Trizol reagent. The extracted RNA was quantified using NanoDrop (Thermo Fisher Scientific, USA). 1 μg of total RNA was used to prepare cDNA using SYBR Green PCR master mix of buffer, random hexamer primers, dNTPs, RNase inhibitors, and reverse transcriptase. The levels were quantitated by the ΔΔC_t_ [47] method using gene-specific primers (Supplementary Table 4) with *RDN18* as the internal control.

### Chromatin Immunoprecipitation (ChIP)

Primary culture was grown in 10 ml YPD media at 30°C for 16 h. Subsequently, 0.1 O.D_.600 nm_ cells from the primary culture were added to fresh culture media and incubated at 30°C, 220 rpm till 1.0 O.D._600 nm_ was reached. Cells were fixed with formaldehyde (1% final concentration) for 15 min. Subsequently, a 10 min treatment with glycine (25 mM) was given to quench formaldehyde. The cells were collected by centrifugation, washed with PBS, and treated with 5 U lyticase enzyme for 2 h. The cells were collected by centrifugation (5000 rpm, 5 min at room temperature), washed, and lysed in lysis buffer (50 mM Tris-Cl, pH 7.5; 140 mM NaCl; 1 mM EDTA; 1% Triton X-100; 1 mM PMSF) with glass beads and vortexed for 5 cycles (30 s ON; 30 s OFF). The supernatant was collected by centrifugation (5000 rpm, 5 min at room temperature) and sonicated in a water bath sonicator (30 s ON; 30 s OFF for 5 cycles). The sheared chromatin was incubated at 4°C for 16 h with the respective antibodies, followed by incubation 4°C for 3 h with Protein G agarose beads. The samples were washed with lysis buffer followed by a wash with high-salt buffer (lysis buffer containing 500 mM NaCl). The samples were washed with wash buffer (10 mM Tris-Cl, pH 8.0; 250 mM LiCl; 1 mM EDTA; 0.5% NP-40), and 50 mM Tris-EDTA pH 8.0 buffer. The chromatin-protein complex was eluted using elution buffer (1.0% SDS; 100 mM NaHCO_3_). Proteinase K (1 μl of 20 mg/ml at 55°C followed by inactivation at 95°C for 10 min) was used to digest the protein in the ChIP complex. The DNA was extracted using 10 % Chelax (Bio-Rad, USA) beads. The ChIP samples were analyzed by qPCR using promoter-specific primers (Supplementary Table 5).

### Growth rate analysis

Primary culture was grown in 10 ml YPD media at 30°C for 16 h. Subsequently, 0.1 O.D_.600 nm_ cells from the primary culture were added to fresh culture media and incubated at 30°C, 220 rpm. The O.D._600 nm_ was monitored every 2 h for 24 h. The doubling time was calculated using the exponential phase data points.

### Drug susceptibility assay

Primary culture was grown in 10 ml YPD media at 30°C for 16 h. Subsequently, 0.1 O.D_.600 nm_ cells from the primary culture were added to fresh culture media and incubated at 30°C, 220 rpm. The secondary culture was grown to 1 O.D._600 nm._ Dilutions were made with 0.9% saline starting from 0.7 O.D._600nm_ and serially diluted 5 times. 5 µl of each dilution were put on plates. YPD plates were used as control. The plates were allowed to dry and then incubated at 30°C for 48 h before photographing.

### Minimal Inhibitory Concentration 80 (MIC80)

Primary culture was grown in 5 ml YPD media at 30°C for 15 h. Subsequently, 0.2 O.D_.600nm_ cells from the primary culture was added to fresh 10 ml culture media and incubated at 30°C, 220 rpm.

Cells from secondary culture were grown to 1 O.D._600nm._ In a 96-well microtiter plate, 100 µl YPD media was added from 2^nd^ to 11^th^ well. A volume of 200 µl sterile media was added to the 12^th^ well, which was taken as a negative control and did not contain any cells. In the first well, 200 µl of the drug was added, and then 2-fold serial dilution was performed by adding 100 µl containing YPD media from one well to the next up to the 10th well. The 11^th^ well was the positive control and therefore, did not have any drug. The cells were resuspended in 0.9% saline to an O.D._600nm_ of 0.1 and added to the wells to a final O.D. _600 nm_ of 0.01. The microtiter plate was incubated at 30°C, 80 rpm for 48 h and then O.D._600 nm_ was measured using a multi-plate reader (Bio-Tek Cytation 5). The MIC80 was calculated at the concentration of the drug where 80% of inhibition of growth was observed [48].

### Rhodamine 6G uptake and efflux assay

Secondary culture grown till 1 O.D._600nm_ was pelleted, washed with PBS and weighed. Equal weight of both the wild type and the mutant strains were taken and incubated with de-energization buffer consisting of 5 mM 2,4-DNP and 5 mM deoxyglucose for 2 h. The cells were then washed again with PBS and incubated with R6G (10 µM) for 1 h. in every 10 min 500 µl of the culture was taken centrifuged and 200 µl of the supernatant was transferred in 96 well plate and fluorescence were read using excitation wavelength of 527 nm; emission wavelength of 555 nm (BioTek Cytation 5).

The R6G efflux assay was performed by the energy-dependent efflux method. Briefly, cells from an overnight culture were inoculated in fresh YEPD medium at 0.1 OD and grown for 4-5 h at 30°C until the log phase. The cell suspension was washed with 1X PBS (phosphate buffered saline) twice and incubated for 3 h at 200 rpm and 30°C for starvation (glucose-free) to reduce the activity of the ABC multi-drug transporters. After incubation, cells were washed twice with PBS and diluted to obtain 10^8^ cells/mL in PBS. R6G at a final concentration of 10 µM was added to the suspension and incubated for 3 h at 30°C and 200 rpm for accumulation assay. For the efflux assay, the cells were washed twice in PBS, 2% glucose was added to the suspension, and incubated for 45 min. The supernatant was then collected, followed by measurement of fluorescence of R6G in a fluorescence spectrophotometer at excitation and emission wavelengths of 527 nm and 555 nm, respectively (Biotek Cytation 5).

### Detection of β-1,3-Glucan and Chitin using confocal microscope

A primary culture of the strains was grown by inoculating a single colony into YPD medium and incubated overnight. Secondary cultures were prepared by inoculating 1% (v/v) of the primary culture into fresh YPD medium and grown to an optical density (O.D._600nm_) of 0.6. The cells were harvested by centrifugation at 5000 rpm for 10 min at 4°C, and the supernatant was discarded. The pellet was washed twice with 1X phosphate-buffered saline (PBS) and subsequently resuspended in 1X PBS containing 100 µg/ml of either Calcofluor White (CFW) or Congo Red (CR). The cells were incubated for 30 min at room temperature and then washed twice with 1X PBS. Finally, the pellet was resuspended in 100 µl of 80% glycerol in 1X PBS, and 10 µl of this suspension was used to prepare slides. The staining was examined using a Confocal microscope (Nikon Eclipse Ti2 Laser Scanning micro-scope) under a 60X oil immersion objective. Quantitation was done using the “intensity line profile” feature of the NIS-Elements AR (Advanced Research) software. Briefly, a circle was drawn around each cell, and the software showed the pixels along this circle. The numbers were individually averaged for all the cells and plotted using GraphPad Prism.

### Statistical analysis

All qPCR and ChIP experiments are reported as the average ± standard error of mean (SEM) of three independent (biological) experiments unless otherwise specified. Each independent experiment was performed as at least two technical replicates. The statistical significance (p-value) was calculated using a paired t-test available in Sigma Plot or Graph Pad prism. The differences were considered significant at p < 0.05.

## Author information

### Author Contributions

Conceptualization, R.M.; Methodology, R.M., and S.B.; Investigation, S.B., A.A., K.P., R.K., and A.G.; Writing—original draft, R.M., and S.B.; Writing—review and editing, R.M., and S.B., Funding acquisition, R.M.; Supervision, R.M. All authors have read and agreed to the published version of the manuscript.

### Funding

The work was supported by CSIR-ASPIRE (37WS (0010)/2023-24/EMR-II/ASPIRE). We acknowledge the facilities/laboratories supported by DBT BUILDER (BT/INF/22/SP45382/2022) and DST FIST-II (SR/FST/LSII-046/2016(C)).

S.B. and R.K. were supported by fellowship from the University Grants Commission, India.

A.G. was supported by UGC Non-Net fellowship.

### Conflict of Interest Statement

The authors report there are no competing interests to declare

### Data Availability Statement

The authors confirm that the data supporting the findings of this study are available within the article and its supplementary materials.

## Notes

### Competing Interest Statement

The authors have declared no competing interest.

