## Supplementary figures and images for "*Candida albicans* Snf2 modulates the response to DNA damage by regulating gene expression and uptake of the genotoxic stressors"

### Fig. S1

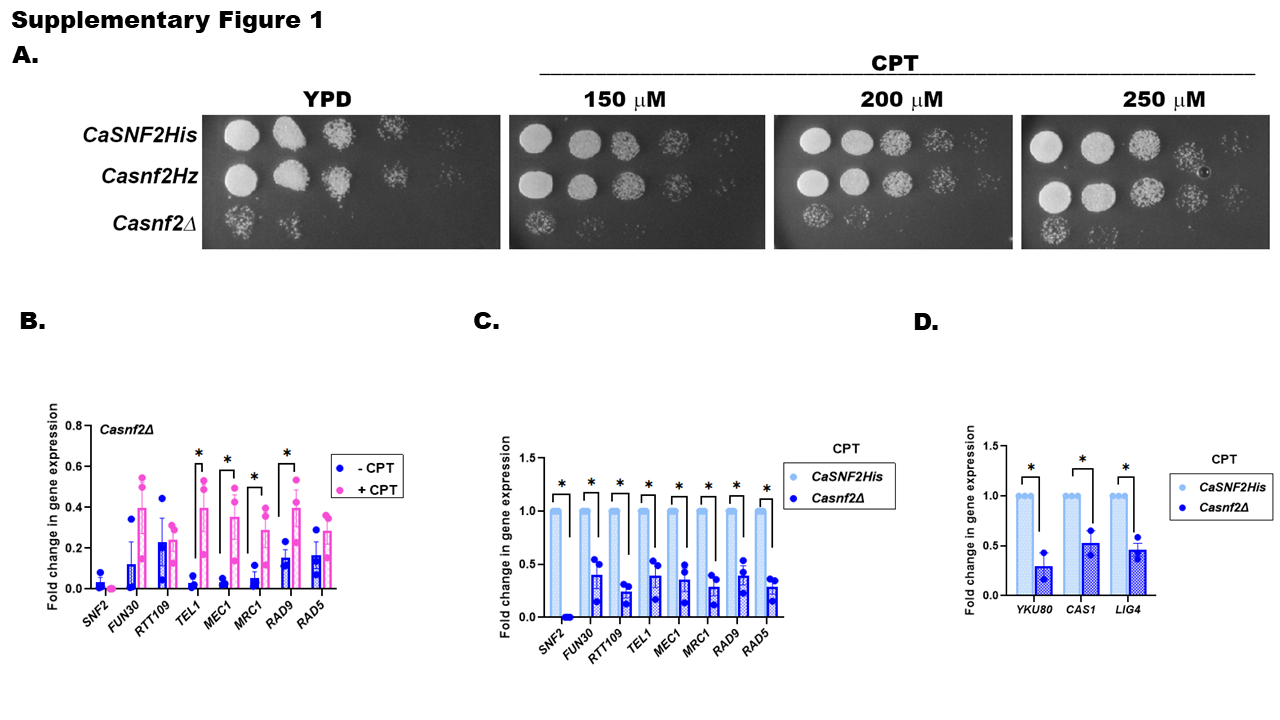

### Fig. S2

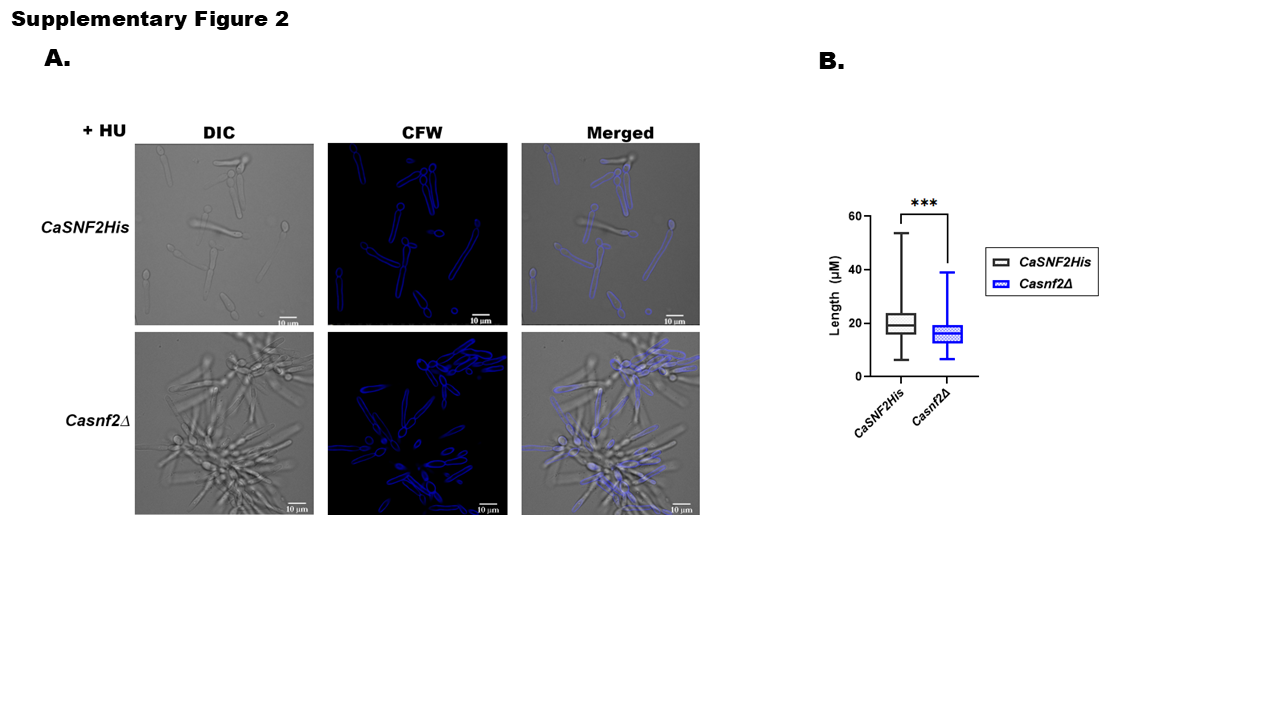

### Fig. S3

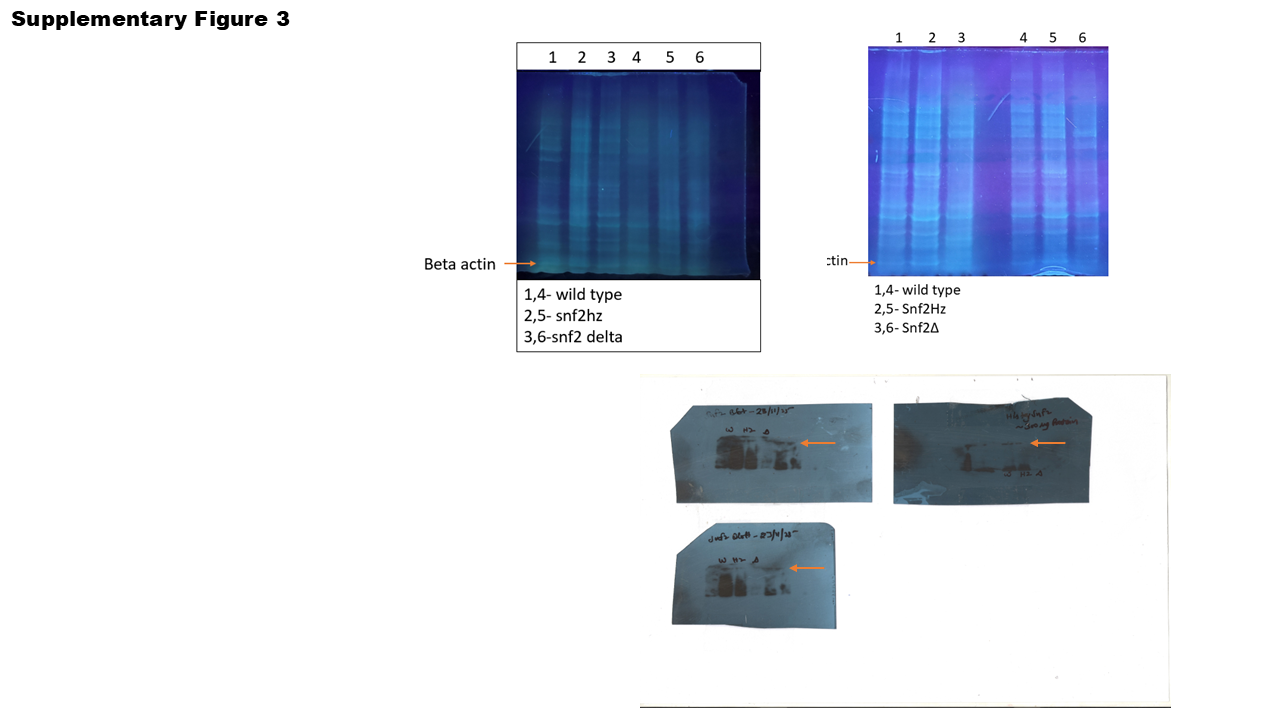
