## Supplementary File for "*Candida albicans* Snf2 modulates the response to DNA damage by regulating gene expression and uptake of the genotoxic stressors"

**Running title:** *Candida albicans* Snf2 regulates the uptake of genotoxic stressors

**Keywords:** *Candida albicans*; ATP-dependent chromatin remodelling; Snf2; DNA damage response;

**SUPPLEMENTARY FIGURE LEGENDS**

**Supplementary Figure 1. *CaSNF2* is not required for CPT-mediated DNA damage response.** (A). Plate assays showing the response of *CaSNF2His* and mutant strains to CPT. (B). Expression of *CaSNF2, CaFUN30, CaRTT109, CaTEL1, CaMEC1, CaMRC1, CaRAD9,* and *CaRAD5* were analyzed in the presence of MMS in *Casnf2Δ* cells in the presence and absence of CPT. (C). Expression of *CaSNF2, CaFUN30, CaRTT109, CaTEL1, CaMEC1, CaMRC1, CaRAD9,* and *CaRAD5* were analyzed in the presence of MMS in *CaSNF2His* and *Casnf2Δ* cells. (D). Expression of *CaYKU80, CaCAS1,* and *CaLIG4* were analyzed in the presence of MMS in *CaSNF2His* and *Casnf2Δ* cells.

**Supplementary Figure 2. Resistance to HU in *Casnf2Δ* mutant is due to cell wall perturbations.** (A). CFW staining of *CaSNF2His* and *Casnf2Δ* cells in the presence of HU. (B) Quantitation of the CFW fluorescence intensity.

**Supplementary Table 1.** List of primers used for creating mutants

| **Primer name** | **Primer sequence (5'-3')** |
| --- | --- |
| **Marker amplification primers*** | |
| Universal P2 | CCGCTGCTAGGCGCGCCGTGACCAGTGTGATGGATATCTGC |
| Universal P5** | GCAGGGATGCGGCCGCTGACAGCTCGGATCCACTAGTAACG |
| ***SNF2* 6X His tag primers** | |
| SNF2 P1 his tag | CTTGACGTGATTAAAAAGAGAACAGC |
| SNF2 P3 his tag | TCAGTGGTGGTGGTGGTGGTGATCAAAATTTGCTGGTGTAGACTC |
| SNF2 P4 his tag | GTCAGCGGCCGCATCCCTGCACAGCATCATTAAGGGTTTTATTCA |
| SNF2 P6 his tag | AACATCATCATCAACCACAAATCCT |
| His tag P2*** | CACCACCACCACCACCACTGAACCAGTGTGATGGATATCTGC |
| His tag check FP3 | CAGAAAGGTGGTACCGACGAAG |
| ***FUN30* myc tag primers** | |
| FUN30 myc tag P1 | GTTCTTGGACAGTGGTAAGGT |
| FUN30 myc tagP3 | TCACAGATCCTCTTCTGAGATGAGTTTTTGTTCACTATAAACTATTGACTCTA |
| FUN30 myc tag P4 | GTCAGCGGCCGCATCCCTGCAGCTGATTTTCCCGCTTGTT |
| FUN30 myc tag P6 | TGTCAATCCACCACCACCAA |
| Myc tag check FP | AGGCAGCATTACATCCGTTG |
| Myc tag P2*** | GAACAAAAACTCATCTCAGAAGAGGATCTGTGAACCAGTGTGATGGATATCTGC |
| ***SNF2* mutant primers** | |
| SNF2 P1 | TTAGCACGTGATTGAACAAAAGAAC |
| SNF2 P3 | CACGGCGCGCCTAGCAGCGGTCCAAACCCAAGGAAAATCTAAAAC |
| SNF2 P4 | GTCAGCGGCCGCATCCCTGCACAGCATCATTAAGGGTTTTATTCA |
| SNF2 P6 | AACATCATCATCAACCACAAATCCT |
| SNF2 del check FP | GGAAATGCCTGTTCGTTTAAGATATT |
| ***RAD9 Hz* mutant primers** | |
| RAD9 P1 | GAGGAGGAGGAGGAGAAAGG |
| RAD9 P3 | CACGGCGCGCCTAGCAGCGGGGCCCCGTCTTACCATATTG |
| RAD9 P4 | GTCAGCGGCCGCATCCCTGCGTTGAGGGTTGGTGTGTCAA |
| RAD9 P6 | GTTGTTGCTTCTCATAATGCTGA |
| RAD9 del check FP | TGGGTGTTTTAAGTGGCCAA |

* Marker amplification Primers are common for every selection marker used in this study i.e., *LEU2, HIS1* and *ARG4*.

**For site specific integration of each cassette Gene del check FP/ Gene tag check (forward primer) and Universal P5 (reverse primer) is used.

*** For marker amplification of each Tag cassette Gene tag P2 (forward primer) and Universal P5 (reverse primer) is used.

**Supplementary Table 2:** List of strains used in this study

| **Strains** | **Genotype** | **Referred to in the manuscript as:** | **Reference** |
| --- | --- | --- | --- |
| SN152 | *ura3/*::imm434::*URA3/ura3*::imm434 *iro1::IRO1/iro1*::imm434*his*1::hisG/*his1*::hisG*leu2/leu2 arg4/arg4* | SN152 | Noble and Johnson,  2005 (24). |
| *SNF2His* | SN152 with *SNF2-*6XHis- *LEU2* | *Casn2His* | This study |
| *SNF2His*/*snf2* | *SNF2His* with *SNF2* /*snf2Δ:: HIS1* | *Casnf2Hz* | This study |
| *snf2/snf2* | SN152 with *snf2Δ:: HIS1* /*snf2 ∆::ARG4* | *Casnf2Δ* | This study |
| *snf2/snf2/ pVT50-SNF2* | *snf2Δ* with *RPS1*/*rps1Δ::pVT50-SNF2* | *Casnf2Δ/SNF2* | This study |
| *snf2/snf2/ pVT50* | *snf2Δ* with *RPS1*/*rps1Δ::pVT50* | *Casnf2Δ /pVT50* | This study |
| *RAD9/rad9* | SN152 with *RAD9/rad9* *:: LEU2* | *Carad9Hz* | This study |
| *snf2/snf2/* *RAD9/rad9* | *snf2Δ* with *RAD9/rad9* *:: LEU2* | *Casnf2Δ /rad9Hz* | This study |
| *MRC1/mrc1* | SN152 with *MRC1/mrc1:: LEU2* | *Camrc1Hz* | This study |
| *snf2/snf2/* *MRC1/mrc1* | *snf2Δ* with *MRC1/mrc1:: LEU2* | *Casnf2Δ/mrc1Hz* | This study |
| *snf2/snf2/ pVT50-MEC1* | *snf2Δ with RPS1/rps1Δ::pVT50-MEC1* | *Casnf2Δ /MEC1* | This study |
| *snf2/snf2/ pVT50-YDC1* | *snf2Δ with RPS1/rps1Δ::pVT50-YDC1* | *Casnf2Δ /YDC1* | This study |
| *snf2/snf2/ pVT50-ACE2* | *snf2Δ with RPS1/rps1Δ::pVT50-ACE2* | *Casnf2Δ /ACE2* | This study |
| *snf2/snf2/ pVT50-RNR1* | *snf2Δ with RPS1/rps1Δ::pVT50-RNR1* | *Casnf2Δ /RNR1* | This study |

**Supplementary Table 3:** List of primers used for creating overexpression and revertant mutants.

| **Primer name** | **Primer sequence (5'-3')** |
| --- | --- |
| ***SNF2* revertant primers** | |
| SNF2 RV FP | CCGCTCGAGCGGATGAATCGTCAACCTACAAGAGAGG |
| SNF2 RV RP | CTAGCTAGCTAGTCAATCAAAATTTGCTGGTGTAGACTCT |
| ***YDC1* overexpression primers** | |
| YDC1 OE FP | CCGCTCGAGCGGATGTTACCTTTTGCCTGGCCATATC |
| YDC1 OE RP | CTAGCTAGCTAGTTAAATGTGTTTCTTATCTTCTTCTGCC |
| ***MEC1* overexpression primers** | |
| MEC1 OE FP | CCGCTCGAGCGGATGACGTCGAATCAATCAATAAGT |
| MEC1 OE RP | CTAGCTAGCTAGCTACATATAAGCTGCCCAACC |
| ***ACE2* overexpression primers** | |
| ACE2 OE FP | CCGCTCGAGCGGATGCATTGGAAATTTCTGAACTTTCG |
| ACE2 OE RP | CTAGCTAGCTAGCTATTGCAACATTAAAAACTCCTCAGTA |

**Supplementary Table 4:** List of primers used in qPCR experiments.

| **Gene** | **Forward primer (5'**-**3')** | **Reverse primer (5'**-**3')** |
| --- | --- | --- |
| *RTT109* | TCGTTGATTGGATGCTGTAAGG | ACCAGCTTCAACAGGTTCATAA |
| *FUN30* | GTGGAATTGAACCAAGTGTAGCTG | TGAGATTGCCTTCCGTTGTCTC |
| *TEL1* | ATTCTACCAGTTGGCTTGCGA | TCCAAAGTTGTTCTTTCCGGC |
| *MEC1* | GAGACACAGCAAGACCCATTA | CGAGCAACTTGTCATCTTTCAG |
| *SNF2* | TGGAGGAGTATGGTCGTGGT | TCTTCGGCTTGGCTTCCATT |
| *RAD5* | GGCCATACGCATCTCAAACT | CGGCGTTTGGATAATCTTGT |
| *RAD9* | TCAAATTCATTGGTGGCTGA | TTCGTTTTCGTTCTCGTTCA |
| *MRC1* | TGACGAACAAGCCACTCAAG | TCATTTTACGACCACGACGA |
| *PMA1* | CAAGAGATGATACTGCTGCC | TGGCAACCAAGTAACCTCTA |
| *RDN18* | CCACCACCCACAAAATCAA | CGGCACCTTACGAGAAATCA |
| *EFB1* | CCAGCTGCCAAATCTATTGT | TCAACAGCAGCTTGTAAGTC |
| *DUN1* | GGTCATTATGCTGTTGTGAAAG | AAGTGTTGACGGAATAGTTGTT |
| *RNR1* | TGGAATTGAGATTACCATTTGAC | TGGCCAAATCTTGTTTTAATGTA |
| *RNR3* | TTTGAACCTTACACTTCGAATTT | CCCAAACGGTTTTATATAGTTGT |
| *LIG4* | GATTGGATAAAAGTCAAACCAGAG | CATTGACACATCTGTTTTAATCCA |
| *YKU80* | ATTTCACTGAAGAAGCAACATATG | TATCATCTGTGTCACTTAAGTCTG |
| *CAS1* | GTTTAGAGACATTTCCCATTTCAG | TTGGATGAGTTTGTTGTATGAGTA |
| *CDR1* | TGGCCATTTATATTGCTTTAACTG | AGTACTACCCTTTTCAGTGAATTT |
| *CDR2* | TGCTGTTTTCTTCTTAGGAGTTTA | AGTACTACCCTTTTCAGTGAATTT |
| *MDR1* | CAGTTTATATGGGATCAGCAGTAT | ACCAGCAATATTATTAACCAAAGC |
| *YDC1* | GAGGCATTGAATACTACCACTA | TGTCTTGAATTCACTAAACACG |
| *MOB1* | ACCACAAACTCCCTTCGACA | CGTCTTCGTC TCGTGGTAAC |
| *ACE2* | ATCAACTCCGCCAACAACAC | TGCTTTGTGGGAGTGTTTGG |
| *DBF2* | CCCCTCCATTTACTCCCCAA | TCGACCCAGGACTAAATCGT |
| *ERG11* | GTGGTGGTAGACATAGATGT | CCATCAATAGTCCATCTTAAA |

**Supplementary Table 5:** List of primers used in ChIP experiments.

| **Gene** | **Forward primer (5'**-**3')** | **Reverse primer (5'**-**3')** |
| --- | --- | --- |
| *FUN30* | AATCAGTTGTATAGGAAGGAGA T | GCAAAAGTTGGTTATTTGTACT AG |
| *SNF2* | GATGAGGGTGGTTGGAGATAT | GGATGAATACTATGTATTGGGT CG |
| *MEC1* | ATTATTCAAGAGACCCTTAAATC CC | ATTTCTATCCTATTTCTGAGGAG GG |
| *TEL1* | GGTAGAGAGAGTGACAGTATCA ACT | ATATCTGACGTAGACATGATTA CG |
| *RTT109* | CCATCCCAGTTAATTGTTTACCA TCTG | GAATACACCACAATAACAACAC TTGCTAAT |
| *RAD5* | CCAACCTTCACTCTAACAAACAT TTAG | TGGGAGAAGTGGTTGTTTCTTC |
| *RAD9* | AGTGTGCACAACTTGAATGG | GTGAATTCTTTGATGAGGCAGA T |
| *MRC1* | GCAGATCCCAATTTGTAACACT | CGTACGGTTTAGCTTAGAAAGA AA |
| *ACE2* | GATTTCAGGTAGCTTCTTCT | TATGTGACTTTCATTTGGAC |
| *YDC1* | CGAGTTGTTGATAAACCATA | GCAATAGCTATCTTCATTTC |
| *GAPDH* | CAGCTGTTTCAAATCCAGGCT | GTGGTTGAGTGGGTTGGTTG |
